# Landscape of super-enhancers in small cell carcinoma of the ovary, hypercalcemic type and efficacy of targeting with natural product triptolide

**DOI:** 10.1101/2023.09.08.556863

**Authors:** Jessica D. Lang, William Selleck, Shawn Striker, Nicolle A. Hipschman, Rochelle Kofman, Anthony N. Karnezis, Felix K. F. Kommoss, Friedrich Kommoss, Jae Rim Wendt, Salvatore J. Facista, William P. D. Hendricks, Krystal A. Orlando, Patrick Pirrotte, Elizabeth A Raupach, Victoria L. Zismann, Yemin Wang, David G. Huntsman, Bernard E. Weissman, Jeffrey M. Trent

**Author notes:** **Financial support**. This work is supported by the National Institutes of Health (R01CA195670 and R01CA195670-S2 to D.G.H., J.T. and B.W. and K99CA234391 to J.D.L), the Canadian Institute of Health Research (CIHR PJT-462168 to Y.W.), the Terry Fox Research Institute Initiative New Frontiers Program in Cancer (D.G.H.), the Marsha Rivkin Center for Ovarian Cancer Research, the Ovarian Cancer Alliance of Arizona, the Small Cell Ovarian Cancer Foundation, Colleen’s Dream Foundation, and philanthropic support to the TGen Foundation. **Corresponding author** Jessica Diane Lang 1111 Highland Avenue, WIMR 2766, Madison, Wisconsin, 53705. **Conflicts of interest**. No conflicts of interest to report.

## Abstract

**Purpose:** Small cell carcinoma of the ovary-hypercalcemic type (SCCOHT) is a rare form of ovarian cancer affecting young women and girls. SCCOHT is driven by loss of both SWI/SNF ATPases SMARCA4 and SMARCA2, having major effects on enhancer landscapes. Super-enhancers are a distinct subset of enhancer clusters frequently associated with oncogenes in cancer.

**Experimental Design:** SCCOHT cell lines and PDX models were interrogated for super-enhancer landscape with H3K27ac CUT&RUN integrated with RNAseq data for associated oncogene analysis. IHC staining and drug efficacy studies in PDX models demonstrate clinical translatability.

**Results:** Here we discovered key distinctions between SWI/SNF chromatin occupancy following SMARCA4 restoration at enhancer vs. super-enhancer sites and characterized putative oncogene expression driven by super-enhancer activity. SCCOHT super-enhancer target genes were particularly enriched in developmental processes, most notably nervous system development. We found high sensitivity of SCCOHT cell lines to triptolide, a small molecule that targets the XPB subunit of the transcription factor II H (TFIIH) complex, found at super-enhancers. Triptolide inhibits expression of many super-enhancer associated genes, including oncogenes. Notably, SALL4 expression is significantly decreased following short triptolide treatment, and its RNA expression was high in SCCOHT tumors relative to other ovarian cancers. In SCCOHT patient-derived xenograft models, triptolide and its prodrug derivative minnelide are particularly effective in inhibiting tumor growth.

**Conclusions:** These results demonstrate the key oncogenic role of super-enhancer activity following epigenetic dysfunction in SCCOHT, which can be effectively targeted through inhibition of its functional components, such as TFIIH inhibition with triptolide.

**Statement of Translational Relevance:** This work identifies a potential therapeutic strategy for small cell carcinoma of the ovary-hypercalcemic type (SCCOHT), a rare and aggressive ovarian cancer affecting young women and children. This study highlights the role of the loss of SWI/SNF ATPase SMARCA4 in altering super-enhancers to promote high oncogene expression. We discovered that SCCOHT cells exhibited high sensitivity to triptolide, a small molecule derived from Tripterygium wilfordii, which targets the XPB subunit of the transcription factor II H (TFIIH) complex found at super-enhancers. Triptolide inhibits the expression of super-enhancer-associated genes, including oncogenes like SALL4, which is highly expressed in SCCOHT. Moreover, in SCCOHT patient-derived xenograft models, triptolide and its derivative minnelide effectively inhibited tumor growth. These findings suggest that targeting super-enhancer activity could be a promising therapeutic approach for SCCOHT, offering potential clinical benefits to patients who currently face limited treatment options and poor outcomes.

## Introduction

Small cell carcinoma of the ovary, hypercalcemic type (SCCOHT) is a rare and aggressive subtype of ovarian cancer that affects women at a mean of 24 years of age. Survival of SCCOHT remains poor with a five-year overall survival rate estimated at 55%, 40%, 29%, and 0% for patients diagnosed at stage I, II, III, and IV, respectively (1). Two-thirds of patients are diagnosed at stage II-IV, necessitating better treatments to improve the survival of these patients with advanced disease. To date, treatment is based on multiple case reports with small patient cohorts. Standard treatment includes multi-agent chemotherapy and radiation, and treatment refractory or relapsed patients have been treated with investigational agents (2).

SCCOHTs are characterized by inactivating germline and/or somatic mutations in *SMARCA4* and concomitant protein loss in over 95% of tumors (3–7). SCCOHTs have low mutational burdens and lack other mutational genetic driver events (3,5,6,8). The loss of SMARCA4 (BRG1) and SMARCA2 (BRM) protein is pathognomonic for SCCOHT amongst other gynecologic cancers and histopathologic mimics (4). SMARCA4 and SMARCA2 are the two mutually exclusive ATPases of the SWItch/Sucrose Non-Fermentable (SWI/SNF) chromatin remodeling complex whose enzymatic activities are necessary for energy-driven histone repositioning throughout the genome. Loss of SWI/SNF ATPases in SCCOHT has been shown to create dependencies on other epigenetic targets, such as polycomb repressive complex 2 (PRC2) and histone deacetylases (HDAC) (4,9–11). The SMARCA4 subunit is particularly important for enhancer function (12,13). While the ATPase activity is essential for regulation of almost a third of enhancers in SCCOHT cells, the residual SWI/SNF lacking SMARCA4 in SCCOHT is retained at 19% of enhancer sites (14). The function of residual SWI/SNF at these sites remains poorly characterized.

Unique activity of super-enhancers (SEs) has been implicated in SWI/SNF-mutant rhabdoid tumors, where SMARCB1-deficient SWI/SNF complex remains localized to SEs, but not regular enhancers (15). SEs are large clusters of enhancers that can regulate expression of many genes, driven through recruitment of RNA polymerase II, transcription factor complexes such as transcription factor II H (TFIIH) and Mediator, and histone modifications characteristic of open chromatin. While SEs in normal cells regulate genes important for cell lineage determination (16), SEs in many tumor types regulate oncogenes leading to their constitutive overexpression (17). SCCOHT, ARID1A-deficient clear cell ovarian carcinoma, and SMARCA4/A2-deficient lung cancer cell lines are sensitive to bromodomain inhibitors that target proteins critical for the maintenance and function of SEs (e.g. BRD4 in Mediator complex) (18–21). This suggests that SWI/SNF-deficient cancers may be dependent on SEs for their growth, despite a lack of general enhancer function in SCCOHT. Characterizing SEs in SCCOHT could identify important therapeutic vulnerabilities in SCCOHT.

Here we describe SE landscapes across SCCOHT cell line models, patient-derived xenografts (PDX), and tumors, including annotation of oncogenes associated with SEs. We further evaluated the feasibility of targeting SEs in SCCOHT using triptolide, which exceptionally decreased the growth of cell lines and PDX models. Triptolide is a naturally-derived drug known to target xeroderma pigmentosum type B (XPB) (22). XPB is an ATP-dependent DNA helicase that functions as part of the TFIIH transcription factor complex essential for assembly of transcriptional machinery and found at high levels at SEs (23). Ultimately, triptolide and SE inhibition may be an effective therapeutic strategy for SCCOHT patients.

## Materials and Methods

### Cell lines

SCCOHT cell lines BIN67 and SCCOHT-1 were maintained in RPMI 1640 (Thermo Fisher Scientific) supplemented with 10% Fetal Bovine Serum (FBS; Thermo Fisher Scientific) and 1% Penicillin/Streptomycin (Thermo Fisher Scientific). COV434 cells were cultured in DMEM (Thermo Fisher Scientific) supplemented with 10% FBS and 1% Penicillin/Streptomycin. COV434 was previously identified as derived from a juvenile granulosa cell tumor, but has now been re-categorized as SCCOHT based on SMARCA4 mutation, lack of SMARCA2 expression and re-evaluation of original tumor specimen (24). All cells were maintained at 37°C in a humidified incubator containing 5% CO_2_ and were routinely monitored for mycoplasma testing and STR profiled for cell line verification.

### Drug Dose Response Assay

SCCOHT cell lines BIN67, SCCOHT-1, and COV434 were each plated into 384-well plates at 1,000 cells per well and incubated at 37°C overnight. A 20-point, 3-fold serial dilution of triptolide ranging from 0.04 nM to 50 μM was applied to cells for 72 hours. CellTiter-Glo (Promega; Cat no. G753) was used per manufacturer’s instructions to determine cell viability and read on a luminescent plate reader. Background-subtracted luminescent readings were analyzed in GraphPad Prism with non-linear regression fit to calculate an IC_50_ value.

### Western blotting

Whole-cell extracts from cell lines were prepared using RIPA buffer (Santa Cruz Biotechnology) with protease and phosphatase inhibitors or 9M urea buffer (9M urea, 4% CHAPS, 0.5% IPG buffer, 50mM DTT, all from Sigma Aldrich) using standard protocols. Thirty µg protein was loaded per well on NuPage 4-12% Bis-Tris gels or Tris Acetate gels (SALL4) and subsequently transferred to PVDF membranes.

For blots detected by chemiluminescence: Blots were pre-blocked in 5% non-fat dry milk in TBST or BSA in TBST for 1 hour and probed using primary antibody overnight. Blots were incubated with secondary antibody (anti-rabbit IgG-HRP, Cell Signaling Technology; anti-mouse IgG-HRP, Santa Cruz Biotechnology) at 1:5000 for 2 hours and developed using Pierce ECL Western Blotting Substrate or SuperSignal West Femto Substrate (Thermo Fisher Scientific).

For blots detected by fluorescence: Blots were blocked in Li-Cor Odyssey Blocking Buffer for 1 hour and probed using primary antibody overnight. Blots were incubated with secondary antibody (IRDye 800CW Goat anti-Mouse #925-32210 or anti-Rabbit #926-32211 and IRDye 680RD Goat anti-Rabbit #925-68071 or anti-Mouse #926-68070, Li-Cor) at 1:20,000 for 2 hours and scanned on Li-Cor CLx.

Primary antibodies: BRD4 (Active Motif, #39909), RPB1 (Cell Signaling Technology, #2629), c-MYC (Cell Signaling Technology, #2276), CyclinD3 (Cell Signaling Technology, #2935), PIM1 (Cell Signaling Technology, #3247), CDK6 (Cell Signaling Technology, #3136), SALL4 (Abcam, ab29112), beta-actin (Cell Signaling Technology, #4970; or ThermoFisher Scientific MA5-15739).

### Animal studies

All procedures were carried out under the institutional guidelines of TGen Drug Development’s Institutional Animal Care and Use Committee (IACUC protocol #19065). Histologically confirmed SMARCA4-mutant SCCOHT tumors PDX-465 and PDX-040 were acquired from Molecular Response (MRL) and serially passaged in mice (4). Tumor suspension in 50% Matrigel / 50% Media in a final volume of 100 μL were subcutaneously inoculated into the backs of NOG (NOD.Cg-*Prkdc^scid^ Il2rg^tm1Sug^*/JicTac) mice. Mice were randomized to treatment arms (n=10) once the average tumor volume reached 75-125 mm^3^. Triptolide (Cayman Chemical) was formulated in DMSO and minnelide (Minneamrita Therapeutics) was formulated in PBS. Vehicle (DMSO or PBS), triptolide (0.6 mg/kg QD days 1-5, QOD days 10-60), or minnelide (0.42 mg/kg QD days 1-26) were administered by intraperitoneal injection (i.p.). Tumor size and body weight were measured twice weekly until endpoint. Tumors were excised upon necropsy and either frozen or formalin-fixed and paraffin-embedded (FFPE).

### Human Specimens

Frozen and FFPE human SCCOHT specimens were obtained under TGen’s Western IRB protocol #1119451, University of North Carolina-Chapel Hill IRB protocol #90-0573, and University of British Columbia Cancer Agency REB project #H19-02823. For the TMA, the ethics committee waived patient consent and approved use of tissue samples (University of Heidelberg, S-463/19). All samples were deidentified and established to be non-human subjects research. Due to the large variance in consenting amongst the SCCOHT patients, clinical information associated with these patients will only be available on the patients eligible for uploading to dbGaP (accession phs001528.v2.p1).

### Tissue microarrays and Immunohistochemistry

Formalin-fixed paraffin-embedded tumor material from the tissue microarray (TMA) with 14 SCCOHT was stained using a mouse monoclonal SALL4 antibody (clone 6E3; RTU; Cell Marque, Rocklin, CA, USA) on a Ventana BenchMark XT immunostainer (Ventana Medical Systems, Tucson, AZ, USA) using standard protocols. SALL4 IHC scoring was performed according to the 3-tier system used by McCluggage et al (25). In brief, a score of 0 indicates no staining, 1 indicates <50% staining, and 2 indicates >50% staining.

### Sequencing preparation and analysis

Details on sequencing can be found in the Supplemental Methods.

### DepMap CRISPR analysis

Publicly available CRISPR data from the DepMap’s Project Achilles (Public 22Q4) were downloaded for BIN67, SCCOHT-1, and COV434 and the 30 identified SE-associated oncogenes from our independent SE analysis. The average Chronos score of the three cell lines was used for plotting against the average change in gene expression following triptolide treatment (see above).

## Results

### SWI/SNF chromatin occupancy at enhancers vs. super-enhancers

Residual SWI/SNF complex in SMARCA4-deficient SCCOHTs has been shown to have dysregulated chromatin occupancy in a manner that largely depends on ATPase loss (14). This previous study showed differences in SWI/SNF occupancy and subunit composition between transcription start site (TSS)-proximal and -distal sites. ARID2-containing PBAF complexes tend to localize to TSS-proximal and DPF2-containing BAF complexes tend to localize to TSS-distal sites upon SMARCA4 restoration (14). In SMARCB1-deficient rhabdoid tumors, loss of SMARCB1 caused differential effects on residual SWI/SNF occupancy at distal enhancers vs. SEs (15). We sought to determine whether disruption of the SWI/SNF ATPase subunit in SCCOHT similarly causes differential effects at these distinct distal sites by re-examining the data produced by the Kadoch group in a SCCOHT cell line, BIN67 (Fig. 1A and Supplemental Fig. 1). This analysis largely recapitulated the distinct BAF vs. PBAF complex localization at proximal and distal sites upon SMARCA4 re-expression. However, we found that distal sites within SE regions displayed an altogether different composition of SWI/SNF, where residual SWI/SNF occupancy was on average higher, including both DPF2 and ARID2 signal.

**Figure 1.**
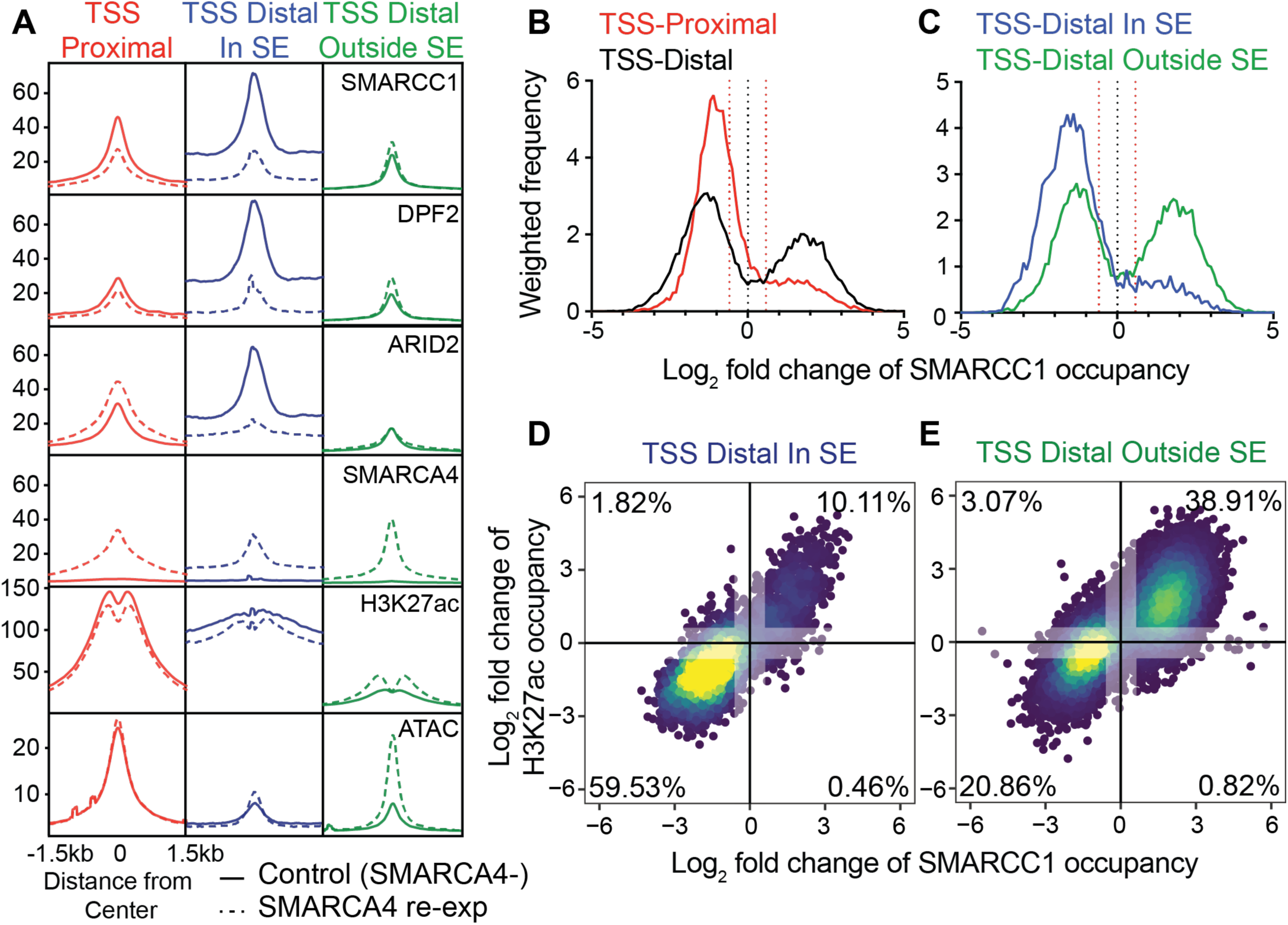
SWI/SNF binding at super-enhancer vs. non-super-enhancer sites. SWI/SNF occupancy sites from BIN67 cells (ChIP data re-analyzed from Pan *et al*. (14) are annotated as proximal (overlapping TSS, 17,866 sites) or distal (>2kb; 18,997 sites) to TSS. Distal sites are separated by those overlapping SE sites (In SE; 4,518 sites), as determined by ROSE on H3K27ac ChIP data, and those non-overlapping those sites (Outside SE; 14,479 sites). (A) Average chromatin occupancy profile of SWI/SNF subunits SMARCC1, DPF2, ARID2, and SMARCA4, as well as H3K27ac and ATAC signal at SWI/SNF-occupied sites, with and without SMARCA4 re-expression. Average ChIP/ATAC signal across region type spanning 1.5kb up- and downstream of peak center is shown. (B & C) Histograms of log_2_ fold-change of SMARCC1 IP signal in SMARCA4 re-expression vs. control expression BIN67 cells at (B) proximal vs. distal SWI/SNF binding sites or (C) distal sites within or outside SE, as described in (A). Histogram frequencies are weighted based on total number of sites in each group (eg. number of proximal vs. distal SWI/SNF binding sites). Positive values indicate more SMARCC1 occupancy at sites defined as SWI/SNF occupied sites from either control or SMARCA4-re-expression conditions, and negative values indicate less SMARCC1 occupancy at those sites with SMARCA4 re-expression. +/-1.5-fold change is demarcated by dotted red line. (D & E) Log_2_ fold-change of SMARCC1 occupancy vs. H3K27ac at each SWI/SNFsite. Plot is colored based on density of points, with yellow being the highest density, and blue being the lowest. The percent of total SWI/SNF sites that had +/- 1.5-fold change in SMARCC1 and H3K27ac is annotated, and values between -1.5 to 1.5 for each are grayed out to indicate which values are considered in this calculation.

Interestingly SMARCA4 re-expression led to a decrease in overall SWI/SNF occupancy at SE sites, as opposed to increases in PBAF at TSS-proximal sites and BAF at TSS-distal sites outside SEs. H3K27ac signal was also reduced at sites of SWI/SNF occupancy in SEs upon SMARCA4 re-expression. Interestingly, chromatin accessibility, as measured by ATAC-seq, was slightly increased at these SE sites. However, looking at the distribution of SWI/SNF occupancy at SEs, the regions with decreased H3K27ac occupancy appeared to have modest changes in accessibility compared to SWI/SNF occupancy sites with increases in H3K27ac, which displayed great increases in accessibility following SMARCA4 re-expression (Supplemental Fig. 1). In fact, three times more SWI/SNF occupancy sites within SEs had decreases in H3K27ac of > 2-fold than SWI/SNF occupancy sites in proximal or distal non-SE sites.

Interestingly, SWI/SNF sites generally showed changes in the core subunit SMARCC1 occupancy with SMARCA4 re-expression (Fig. 1B & 1C). Distal sites more of an increase in SMARCC1 occupancy after SMARCA4 re-expression than proximal sites (Fig. 1B). However, distal sites within SEs displayed even stronger decreases in SMARCC1 occupancy, with only a minority of sites displaying increased SMARCC1 occupancy (Fig. 1C). In addition, upon SMARCA4 re-expression, SMARCC1 occupancy changes correlated with H3K27ac occupancy changes across distal SWI/SNF sites (Fig. 1D & 1E). Distal sites in SEs, however, were more likely to have concomitant decreases in SMARCC1 and H3K27ac across sites, with 59.53% showing dual decreases in SE sites vs. only 20.86% of non-SE distal sites. Together, these data suggest that the SMARCA4-deficiency creates a unique SE state that may be important for the epigenetic reprogramming in SCCOHT.

### Landscape of super-enhancers in SCCOHT cell lines

To more comprehensively examine SE landscapes in SCCOHT, we performed H3K27ac CUT&RUN in the three available SCCOHT cell lines, BIN67, COV434, and SCCOHT-1. Combining results from two SE callers, ROSE and CREAM, we identified 478 SEs in BIN67, 308 SEs in COV434, and 408 SEs in SCCOHT-1 (Fig. 2A). While the majority of SEs (61.8-71.4%) were unique to each cell line, a total of 176 (28.6-38.2%) of SE regions overlapped in more than one cell line, with 40 regions (8.4-13.0%) overlapping in all three SCCOHT cell lines (Fig. 2B). We examined gene expression from SCCOHT cell lines and tumor RNA-seq of the 401 genes located within 50kb of the 176 SE regions detected in at least 2 SCCOHT cell lines (Fig. 2C).

**Figure 2.**
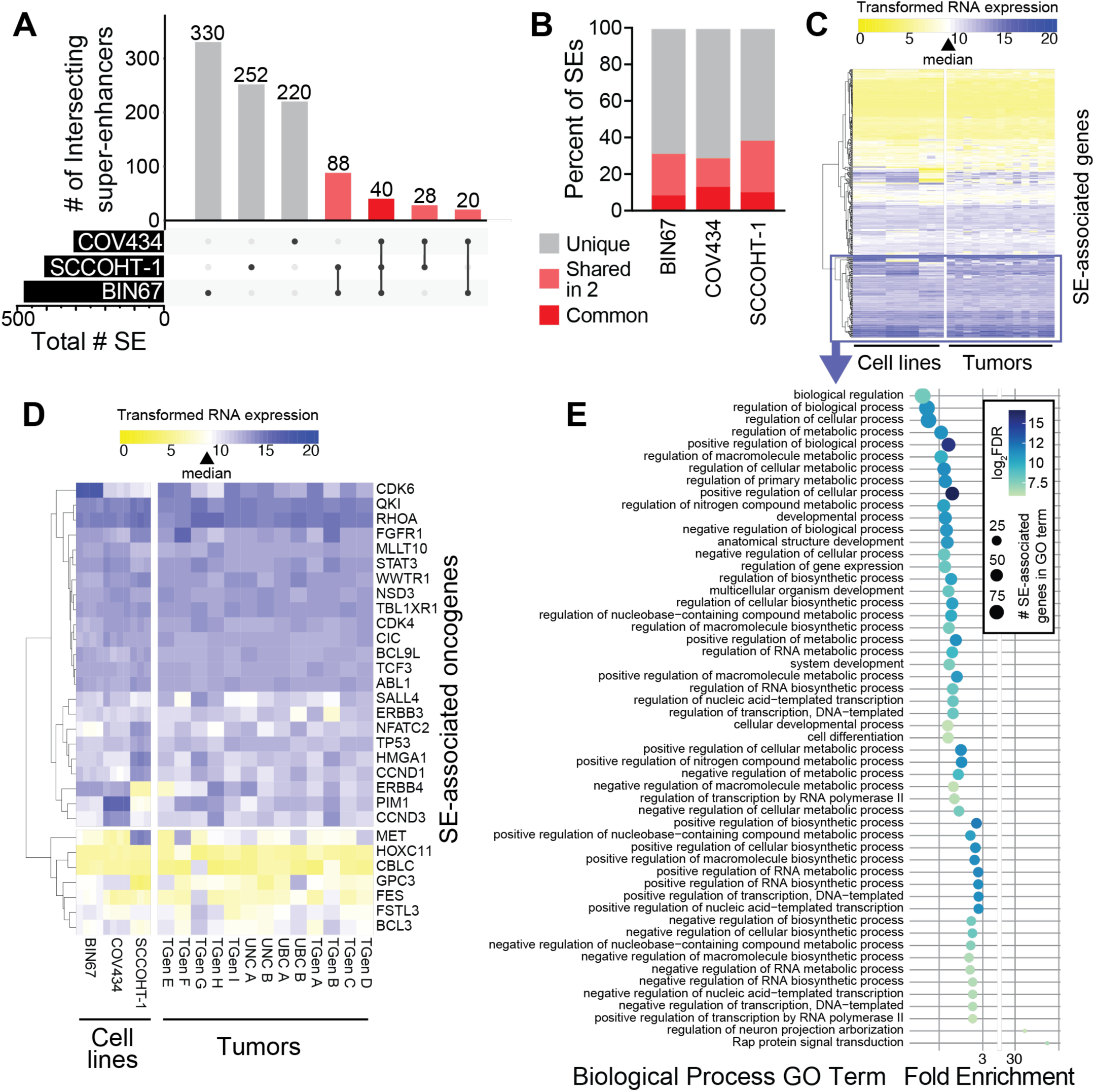
SE landscape and associated gene expression in SCCOHT cell lines and tumors. (A) UpSet plot of overlapping SE regions in three SCCOHT cell lines. SEs were identified by ROSE or CREAM across three replicate H3K27ac CUT&RUN experiments in each cell line were merged for each cell line, then compared using Intervene. Overlapping SE regions are shown in shades of red, with bold red representing overlap in all three cell lines. (B) For each cell line, percentage of SEs classifying as "unique", "shared in 2", or "common" are shown. Colors match those in (A). (C-D) Unsupervised hierarchical clustering of variance stabilizing transformed-TPM RNA levels from RNA-seq of (C) 558 genes within 50kb of the 176 SEs identified in more than one SCCOHT cell line or (D) 30 oncogenes within 50kb of any SE identified in SCCOHT cell lines. Three to four replicates of RNA-seq in SCCOHT cell lines (left) or single RNA-seq analyses from thirteen SCCOHT tumors (right) are shown. Sample order is the same in each panel. The color scale is centered on a white value indicating the median expression value of all protein coding genes across samples (indicated by triangle in legend), such that purple indicates higher expression and yellow indicates lower expression than any other gene annotated in the genome. The top two clusters in (C) with high expression across cell lines and tumors are highlighted with the purple box, and were selected for further gene ontology analysis in (E). (E) Gene ontology (GO) analysis of highly expressed, SE-associated genes using Panther. Bubble plot shows log_2_ enrichment score on the x-axis, log_2_ FDR as bubble color, and size of the SE-associated genes within each term as the bubble size.

We identified 30 oncogenes within proximity to SEs identified in any of the SCCOHT cell lines. 85% of these oncogenes were expressed higher than the protein-coding gene median (Fig. 2D). A notable cluster of oncogenes were highly expressed across all three cell lines and 13 SCCOHT tumors. These included CDK4, CDK6, and FGFR1, which were found previously to be therapeutic targets in SCCOHT models (26,27). ERBB3 and ERBB4 were also high and associated with SEs.

Gene ontology (GO) analyses identified enrichment of developmental and metabolic biological processes among the most highly expressed clusters of SE-associated genes (purple box in Fig. 2C) in SCCOHT (Fig. 2E; Supplemental Table 2). The most statistically significant GO term was "developmental process", which was also related to significant terms "anatomical structure development", "nervous system development", "multicellular organism development", "system development", and "cellular developmental process". This is interesting given roles of SEs in normal cell processes which include lineage specification. Other themes in GO terms included general biologic and metabolic process regulation, as well as more specific processes including "IRE1-mediated unfolded protein response".

### SALL4 and MLLT10 expression is high in SCCOHT

We examined the 30 oncogenes associated with SEs between the 3 SCCOHT cell lines for expression in SCCOHT relative to age-matched TCGA high-grade serous ovarian carcinoma (HGSC) tumors, normal ovary tissue from ENCODE (ENCODE ovary), and ENCODE reference cell lines (ENCODE reference). Patterns of specificity of high expression differed among these 30 genes (Supplemental Fig. 2). SALL4 and MLLT10 were the most selectively expressed in SCCOHT tumors relative to either HGSC or normal samples (Fig. 3A). Consistent with this observation, the SE regions upstream of SALL4 and MLLT10 had consistent high H3K27ac signal in SCCOHT cell lines, but reduced H3K27ac signal across 7 ENCODE reference cell lines (Fig. 3B & C). We also examined SALL4 expression in SCCOHT tumors by IHC. In a SCCOHT TMA with 14 SCCOHT patients, 8 (57%) had positive SALL4 staining (Supplemental Fig. 3). Staining was typically weak and in <20% of the cells in a given area. SALL4 staining was focally strong in one tumor sample.

**Figure 3.**
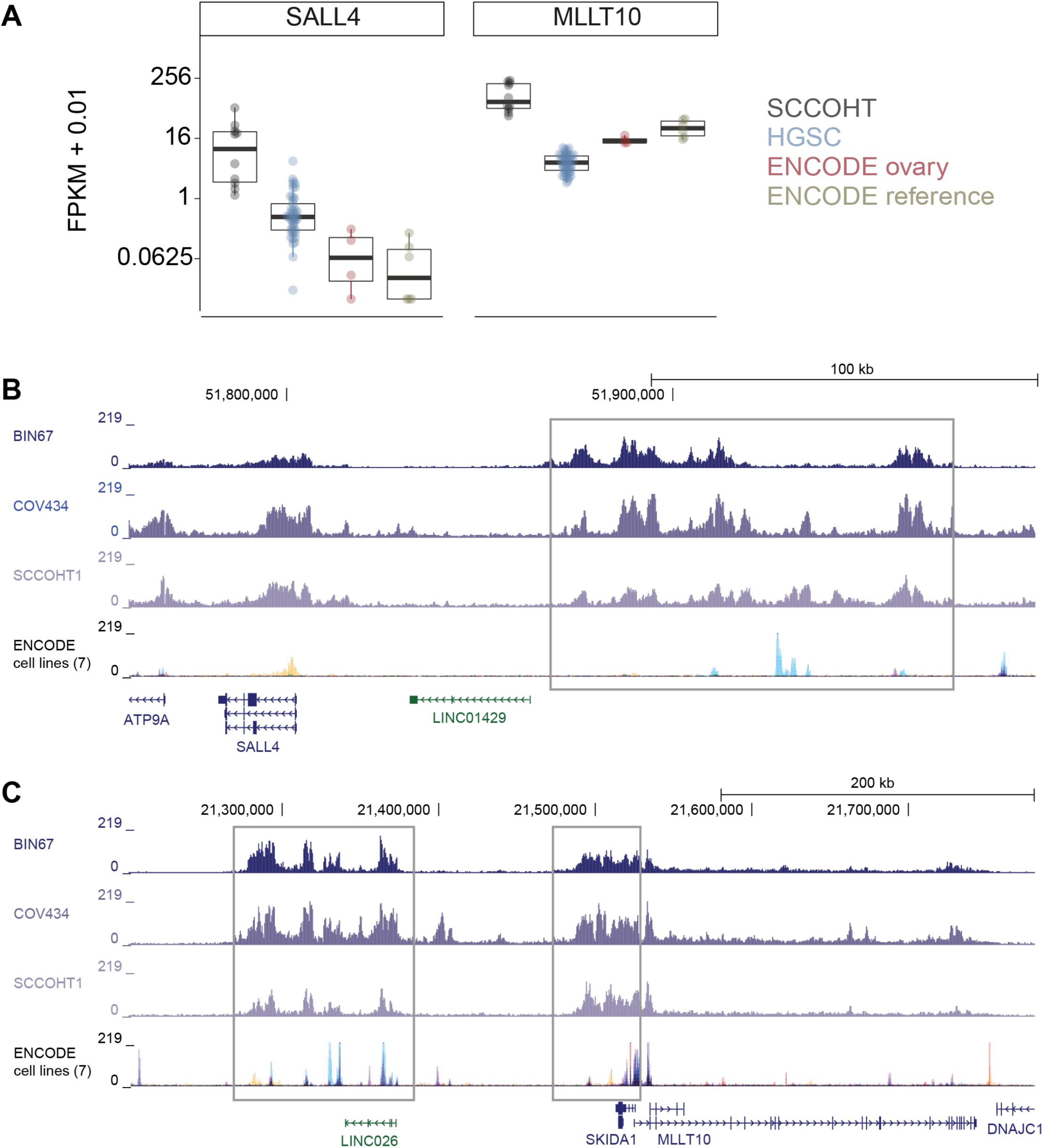
Oncogenes SALL4 and MLLT10 are highly expressed in SCCOHT through unique super-enhancer. (A) RNA expression of SALL4 and MLLT10 from RNA-seq on SCCOHT tumors, age-matched TCGA high-grade serous ovarian carcinoma (HGSC) tumors, normal ovary tissue from ENCODE (ENCODE ovary), and ENCODE reference cell lines (ENCODE reference). FPKM + 0.01 is plotted on a logarithmic scale, with individual circles representing each sample and the box and whiskers plot summarizing the median value (heavy line), upper and lower 25% quartiles (box limits), and error bars showing highest and lowest values. (B & C) H3K27ac tracks at SALL4 (B) and MLLT10 (C) SEs. The gray boxes indicate the common SE regions between the 3 SCCOHT cell lines. The H3K27ac ChIP-seq tracks from the ENCODE reference cell lines (GM12878, H1-hESC, HSMM, HUVEC, K562, NHEK, NHLF) is shown on the bottom track for comparison, with each track overlaid according to UCSC Genome Browser’s default settings and colors.

### Triptolide inhibits super-enhancers in SCCOHT

We next assessed whether we could pharmacologically target SEs in SCCOHT using triptolide, a drug shown to target the XPB subunit of the TFIIH complex (22). Using drug dose response assays, triptolide potently inhibited the growth of all three SCCOHT cell lines, BIN67, SCCOHT-1, and COV434, with IC_50_ values in the low nanomolar range (2-12 nM) (Fig. 4A).

**Figure 4.**
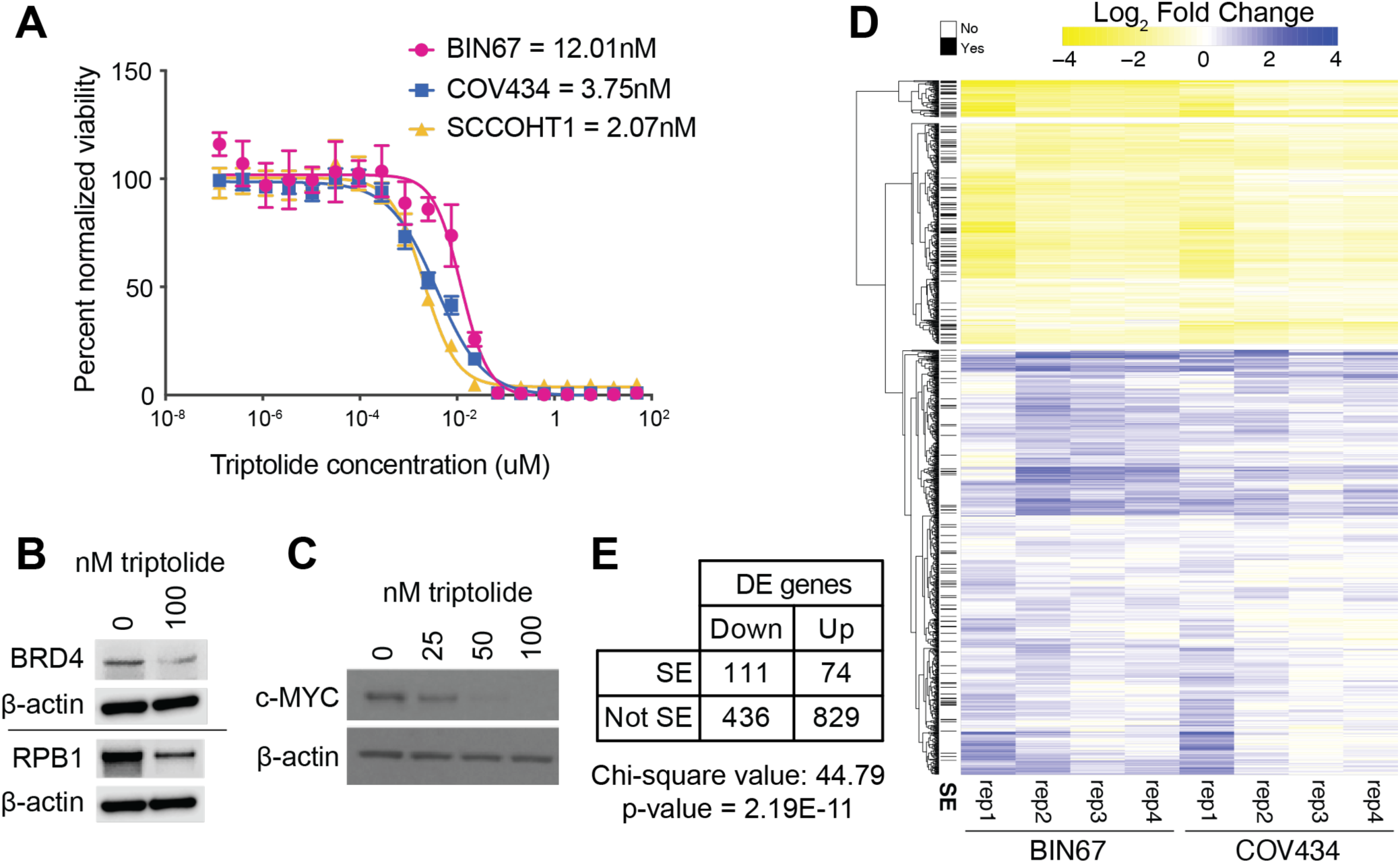
Triptolide inhibits super-enhancers in SCCOHT. (A) Drug dose response assay on SCCOHT cell lines treated with triptolide for 72 hours. IC_50_s for each SCCOHT cell line are provided in the key. (B) BRD4 and RPB1 western blots on BIN67 cells treated with vehicle (0.01% DMSO) or 100 nM Triptolide for 24 hours. β-actin serves as a loading control. (D) c-Myc western blot on BIN67 cells treated with serial dilutions of triptolide for 24 hours. β-actin serves as a loading control. (D) Differential gene expression in SCCOHT cell lines BIN67 and SCCOHT1 after 6 hours treatment with triptolide. Yellow represents a decrease in expression with triptolide treatment, and purple represents an increase. Genes within 50 kb of a SE are denoted on the left with a black line (SE column). (E) Summary of differentially expressed genes from data in (D). DE genes were annotated by direction of gene expression change and whether they were 50kb from a SE. Chi-square test was run to determine whether there was a statistically significant association between direction of change and being near a SE.

Triptolide treatment reduced global protein levels of BRD4 and RPB1 (Fig. 4B), which are important for SE function through their participation in the Mediator and RNA polymerase II complexes, respectively. These global changes are a direct consequence of targeting of XPB by triptolide(22). As one of the most commonly SE-associated oncogenes, c-Myc protein levels decreased in a dose-dependent manner with triptolide treatment (Fig. 4C).

To determine whether triptolide inhibits SEs in SCCOHT cells following treatment, we performed H3K27ac CUT&RUN and RNA-seq in BIN67 and COV434 cells after a six-hour treatment with triptolide. A total of 974 and 460 up-regulated genes and 407 and 155 down-regulated genes were significantly changed following triptolide treatment in BIN67 and COV434 cell lines, respectively, for a total of 903 and 547 up- and down-regulated genes overall (Fig. 4D-E, Fig. 5A-B). The ratio of SE-associated genes that were down-regulated vs. upregulated (1.50) was much higher than for non-SE genes (0.52), which was statistically significant (Fig. 4E).

**Figure 5.**
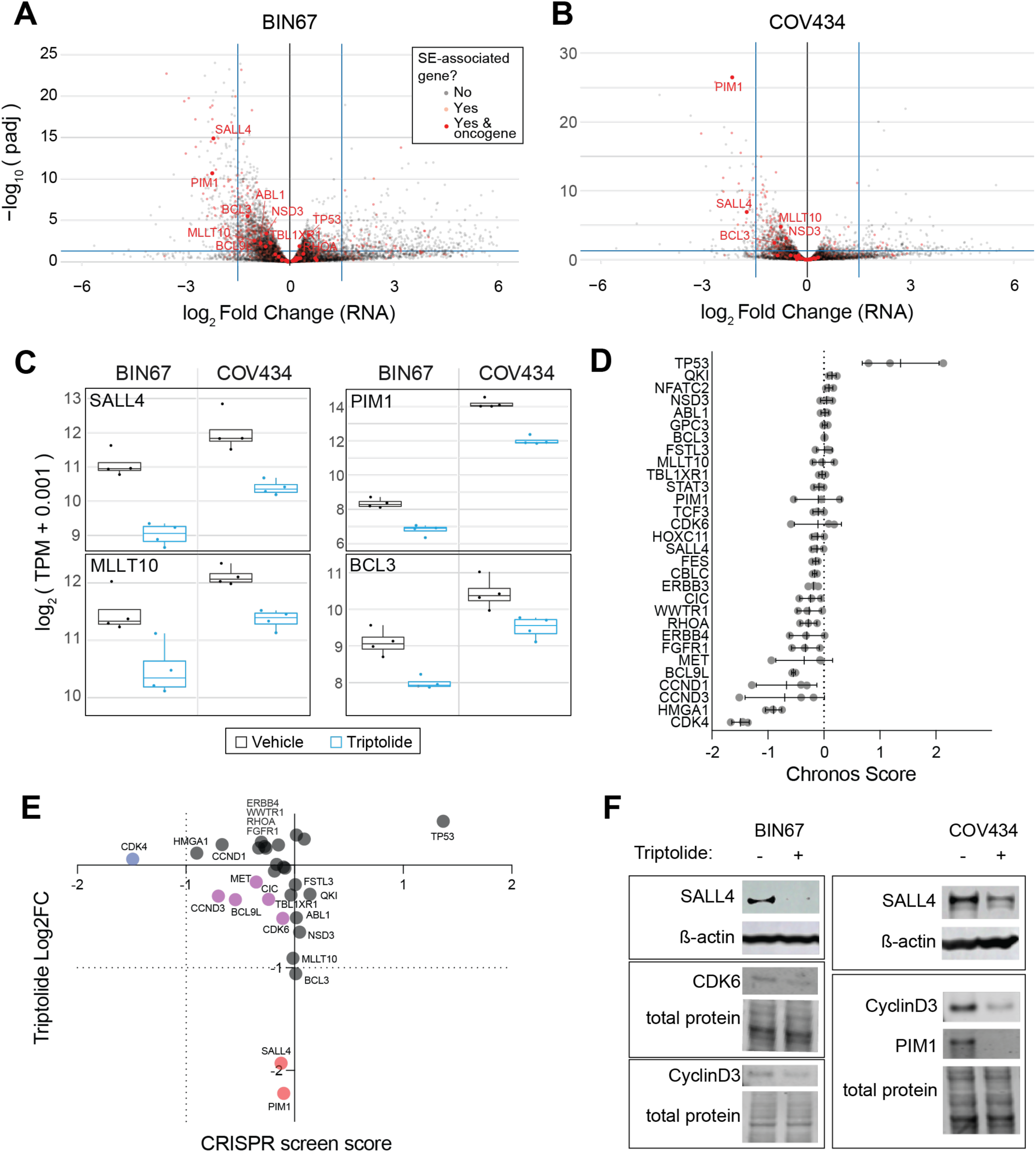
Triptolide inhibits the expression of SE-associated oncogenes. (A & B) Volcano plots of differential gene expression of BIN67 (A) and COV434 (B) after treatment with triptolide for 6 h. Blue bars denote p-value = 0.05 and log_2_ Fold Change = ±1.5. SE-associated genes are denoted by red dots, with non-transparent dots indicating SE-associated oncogenes. Those meeting either cutoff are labeled with gene names. (C) RNA expression of notable SE-associated oncogenes with repression of expression by triptolide. (D) Mean CRISPR screen cell viability score (Chronos score) for SE-associated oncogenes in BIN67, COV434, and SCCOHT1. Error bars represent +/- one standard deviation. (E) SCCOHT cell line average DepMap CRISPR screen score versus avg triptolide treated log2 fold change in RNA expression of SE-associated oncogenes. Gene names are as indicated, with exception of clusters with no appreciable difference in either assay. Genes highlighted in blue indicate CRISPR screen specific hits, red indicate differential expression hits, and purple indicate some intermediate hits on both screens. (F) Western blots for SE-associated oncogenes in BIN67 and COV434 treated with 50 nM triptolide for 24 hours. For SALL4, beta-actin serves as a loading control, and for all other proteins, total protein stain was used as a loading control.

Interestingly, H3K27ac was largely static at the same 6-hour timepoint after triptolide treatment, suggesting that triptolide functions downstream of establishment of the open chromatin environment, consistent with its role in inhibiting TFIIH (Supplemental Fig. 4). Only one peak was significant, increasing only in BIN67, and it was not associated with a SE. By 16 hours, H3K27ac starts to decline at some SEs (Supplemental Fig. 4). COV434 showed fewer down-regulated SE peaks than observed in BIN67, likely due to the overall fewer SEs near oncogenes identified in COV434. In BIN67, H3K27ac peaks within SEs that displayed decreases included those near SALL4, TMSB4X, MLLT10, PIM1, BCL3, CBLC, HOXC11, and MALAT1.

### SALL4, PIM1, and MLLT10 expression is significantly reduced by triptolide

We examined expression of SE-associated oncogenes to address whether inhibition of these oncogenes may be involved in the mechanism of triptolide function. Overall, expression of SE-associated oncogenes either decreased or had minimal change (Fig. 5A-B). In both BIN67 and COV434, PIM1 and SALL4 were the only SE-associated oncogenes with statistically significant decreased expression (Fig. 5A-C). Many of the SE-associated oncogenes displayed a trend towards decreased expression, although most did not pass the threshold of statistical significance (Fig. 5C).

To determine whether SE-associated oncogenes are essential for the growth of SCCOHT tumors, we looked into publicly available genome-wide CRISPR knockout data on the three SCCOHT cell lines. DepMap’s Achilles Project has included the three SCCOHT cell lines in their CRISPR screen data. Based on the average Chronos score for the SE-associated oncogenes, they were more likely to have a detrimental effect on cell growth (Fig. 5D), though the effect size of targeting most single SE-associated genes was not large. Interestingly, TP53 knockout increased growth of SCCOHT cell lines, consistent with its canonical tumor suppressor activity in cancer. We left TP53 in the analysis for two reasons: (1) to remain unbiased, as COSMIC identified it as a potential oncogene in some cancer contexts, and (2) based on specific observations in SMARCB1-deficient rhabdoid tumors which show that TP53 is a synthetic lethal target (28,29). The correlation between RNA expression changes following triptolide treatment and the effect of knockout in SCCOHT cell lines was not linearly associated, with an R-squared value of 0.0022 (Fig. 5E). However, a few genes did have concordant negative effects on both gene expression and growth, including CCND3, BCL9L, MET, CIC, and CDK6.

To confirm down-regulation of the SE-associated oncogenes, protein abundance was examined by western blotting. In BIN67 cells, we confirmed that triptolide reduced SALL4, CDK6, and CyclinD3 protein levels. In COV434, we confirmed down-regulation of SALL4, CyclinD3, and PIM1 (Fig. 5F). PIM1 was not detected in BIN67, and CDK6 was not detected in COV434.

### Triptolide and minnelide are potent in preclinical models of SCCOHT

To determine the preclinical efficacy of triptolide for SCCOHT, we performed *in vivo* efficacy studies in patient-derived xenograft (PDX) models PDX-040 and PDX-465 (27). Triptolide treatment of PDX-040 reduced tumor growth by 77.9% at 30 days of treatment and continued to repress tumor growth through 60 days of treatment (Fig. 6A & B; individual mouse tumor weights in Supplemental Fig. 5A & B). Since triptolide is known to be less bioavailable due to poor water solubility, we also tested the activity of its more water-soluble prodrug, minnelide (30). Minnelide reduced growth of PDX-465 tumors, with an overall reduction of 31.5% in tumor volume since the initiation of treatment, whereas the vehicle-treated PDX-465 tumors had increased in volume of 1,030% (Fig. 6C & D). Overall, mice tolerated both triptolide and minnelide, with mice maintaining stable body weight over the course of treatment (Supplemental Fig. 5C & D). Starting tumor volumes were also equivalent in control and treatment groups, demonstrating that mice were appropriately randomized at initiation of treatment (Supplemental Fig. 5E & F).

**Figure 6.**
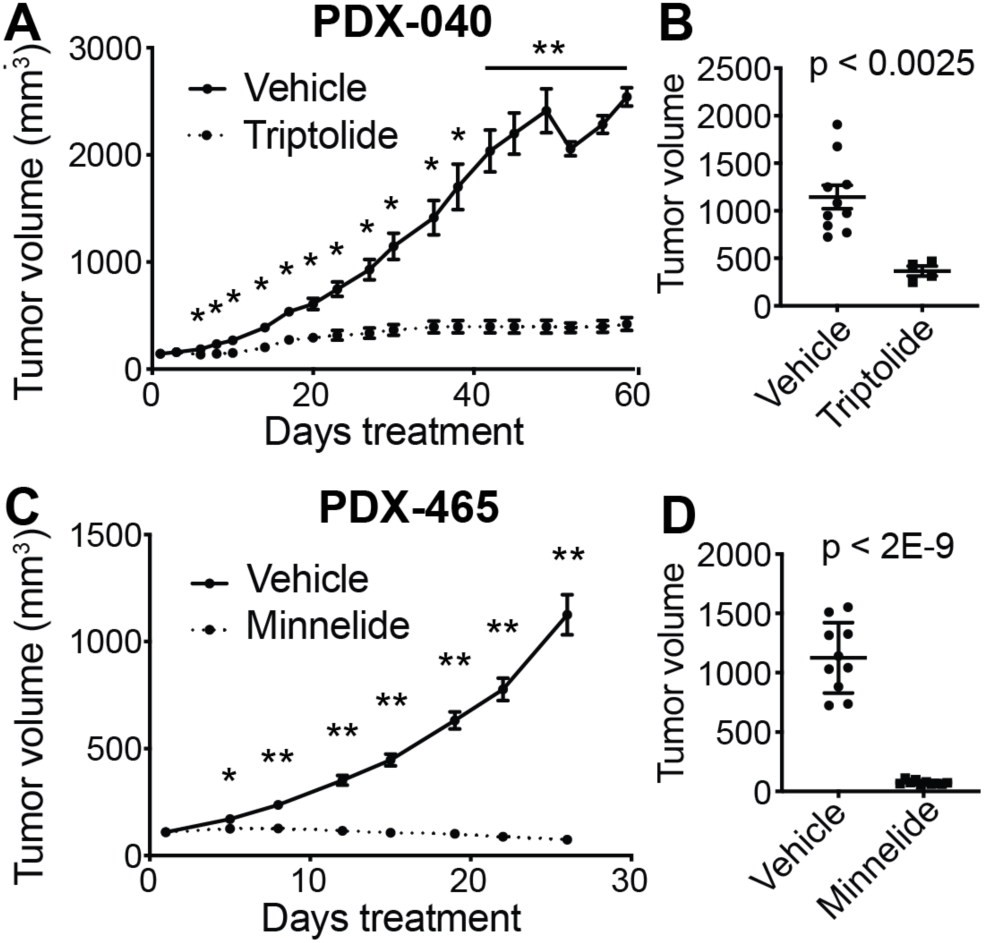
Efficacy of triptolide/minnelide in SCCOHT *in vivo*. (A & B) SCCOHT PDX model 040 treated with vehicle (DMSO i.p.; N=10) or triptolide (0.6 mg/kg i.p., QD days 1-5, QOD days 10-60, N=4). (A) Tumor volume measurements from initiation of treatment (day 1) to study end (day 60). Multiple t-tests were used to determine p-value. * < 0.05, ** < 0.0001. (B) Tumor volume measurements at day 30. Student’s t-test was used to determine p-value. (C & D) SCCOHT PDX model 465 treated with vehicle (PBS i.p., N=10) or minnelide (0.42 mg/kg i.p., QD, N=10). (C) Tumor volume measurements from initiation of treatment (day 1) to study end (day 26). Paired t-test was used to determine p-value. (D) Tumor volume measurements at day 26. Student’s t-test was used to determine p-value. Error bars represent standard error of the mean throughout.

## Discussion

This report is the first to characterize the SE landscape of SCCOHT, an ovarian cancer subtype driven by widespread epigenetic dysregulation due to loss of the SWI/SNF chromatin remodeling complex ATPase subunits SMARCA4 and SMARCA2. We demonstrate that the 30 oncogenes found near SEs detected in more than one SCCOHT cell line are also highly expressed in an independent cohort of SCCOHT tumors. Expression of these oncogenes, such as SALL4 and MLLT10, distinguish SCCOHT tumors from HGSC tumors, normal ovary and other reference cell lines, suggesting their importance broadly in the oncogenic processes of SCCOHT.

SALL4 was associated with a SE in both BIN67 and SCCOHT-1 cells. Given its role as a transcription factor in early embryonic development and maintaining cell stemness, SALL4 may be crucial for maintaining SCCOHT’s undifferentiated phenotype. The low abundance of SALL4 expression in differentiated adult tissues, along with its potential role as an immunogenic antigen (31), could at least partially underlie the perplexing sensitivity of low-mutation burden SCCOHTs to immune checkpoint inhibitors (32). This theory is yet untested in SCCOHT tumors, but is worth exploring in future directions. One immunohistochemistry study has also validated the high expression of SALL4 in SCCOHT compared high-grade serous ovarian cancer (25). Rhabdoid tumors have also been shown to have a SE resident at an overlapping position upstream of the SALL4 gene, and similarly depend upon its expression for rhabdoid tumor cell line growth (15).

In our reanalysis of SCCOHT data with simultaneously prepared SWI/SNF subunit and H3K27ac ChIP-seq and ATAC-seq, we observed that SWI/SNF occupancy sites in SEs differed from those of non-SE distal sites. Following SMARCA4 re-expression, SMARCA4 occupancy increased at SE sites, while other subunits showed overall reductions. This pattern distinct is from the those observed at transcription start sites or other distal binding sites. Additionally, total H3K27ac signal at the SE sites was reduced. Together, this supports a model where SMARCA4 loss in SCCOHT strengthens SE sites, where SMARCA4-deficient SWI/SNF complexes reside despite their lack of ATPase function. Wang *et al*. similarly looked at SWI/SNF complex occupancy at SEs versus other distal sites in SMARCB1-deficient rhabdoid tumors (15). In rhabdoid tumors, SMARCB1 re-expression increased SWI/SNF occupancy at distal sites generally, but not at SE sites, where it remained unchanged. Our data demonstrate a loss of SWI/SNF occupancy at SEs with SMARCA4 re-expression, which is in contrast to what is observed in rhabdoid tumors. In rhabdoid tumors, the loss of SMARCB1 does not interfere with the non-canonical BAF (ncBAF) complex, which requires a functional ATPase subunit.

However, in SCCOHT tumors, the SWI/SNF complex only exists in a residual form distinct and less abundant than ncBAF (33). Further, the rhabdoid tumor data did not show a corresponding loss of H3K27ac signal at SEs. Thus it is highly likely that the underlying differences observed at SEs between SCCOHTs and rhabdoid tumors is driven by the different forms of the SWI/SNF complex predominant based on the specific subunit affected.

Inhibitors can target SEs through essential components of the transcriptional machinery at these sites, such as bromodomain-containing proteins, CDK7, and TFIIH. The exceptional growth inhibition by the SE-targeting small molecule triptolide in SCCOHT models exceeds responses observed by other published inhibitors by us and others (9–11,18,26,27). Among known targets of triptolide is XPB, a subunit of the TFIIH complex. Inhibition of XPB activity may account for the observed decreases in the global expression of its associated transcriptional machinery, including BRD4 and RNAPII subunit RPB1, as well as expression of several oncogenes in close proximity to SEs identified in SCCOHT cell lines. Noel *et al.* observed similar effects of triptolide on H3K27ac at SEs in pancreatic cancer (34). In their co-culture model, they observed more dramatic transcriptional effects to the stromal microenvironment than the tumor cells themselves, yet dramatic effects on cancer growth. Our study did not explore tumor microenvironment effects, so we cannot exclude that this had some effect in our animal models, yet our cell lines alone were quite sensitive.

In our data, we surprisingly see that genes strongly down-regulated by short triptolide treatment were not the same genes that reduced growth in genome-wide CRISPR KO screens performed by DepMap in the same cell lines. However, it is possible that these findings indicate a multi-gene effect, where the loss of SMARCA4 in SCCOHT leads to more SE activity at oncogenes at several sites. This is supported by our analysis of the aforementioned data where re-expression of wild-type SMARCA4 resulted in strongly reduced H3K27ac signal at around two thousand SWI/SNF occupancy sites. It is possible that gene-by-gene screens do not have strong effects on their own; however, simultaneously targeting SEs genome-wide using triptolide has stronger effects.

Recently, Shorstova *et al*. described sensitivity of SMARCA4/A2 dual-deficient cell lines, including the SCCOHT cell line SCCOHT-1, to the bromodomain inhibitors JQ1 and OTX015. Interestingly, they found that ERBB3 is repressed following treatment with OTX015. We found ERBB3 to be in close proximity to SEs in SCCOHT-1 cells, providing a potential mechanism for the effect of bromodomain inhibitors. We also found both CDK4 and CDK6 near SEs in SCCOHT cell lines. We also previously demonstrated sensitivity of SCCOHTs to receptor tyrosine kinase inhibitor, ponatinib (27). FGFR1, one target of ponatinib, is also associated with a SE in SCCOHT cell lines.

The efficacy of SE inhibitors in cancer has been debated based on the potential side effects of targeting general transcription machinery in cancer cells and in normal tissues. The possibility exists that clinically relevant doses of these inhibitors have more specific effects on SEs, as these non-saturating doses likely primarily affect regions of high transcriptional activity. We did not observe general toxicity in our PDX efficacy models after treatment with doses of triptolide and minnelide that resulted in stable disease or tumor regression. Minnelide is being tested in clinical trials for human cancers, with two completed but not yet reported trials in pancreatic (NCT03117920; phase II) and gastrointestinal tumors (NCT01927965; phase I).

Additional trials are also currently recruiting: three phase I, two phase Ib, and a phase II trial in pancreatic cancer. Full reports of these studies have not yet been released. Early phase I results demonstrate reasonable tolerability of the drug, with some cases of hematologic toxicity. Early reports from a pharmacokinetic study of minnelide report two pancreatic cancer patients with some clinical benefit for 7 months (35). Based on our data that demonstrates a unique feature of SE dysregulation following SMARCA4 loss, associated gene expression programs regulating oncogenes, and exceptional responses observed in animal models, triptolide should be considered as a promising treatment for SCCOHT patients.

## Acknowledgements

This work is supported by the National Institutes of Health (R01CA195670 and R01CA195670-S2 to D.G.H., J.T. and B.W. and K99CA234391 to J.D.L), the Canadian Institute of Health Research (CIHR PJT-462168 to Y.W.), the Terry Fox Research Institute Initiative New Frontiers Program in Cancer (D.G.H.), the Marsha Rivkin Center for Ovarian Cancer Research, the Ovarian Cancer Alliance of Arizona, the Small Cell Ovarian Cancer Foundation, Colleen’s Dream Foundation, and philanthropic support to the TGen Foundation. Thank you to Karthigayini Sivaprakasam for her bioinformatics advice. We would also like to thank the SCCOHT patients, their families and communities, and the clinicians who have contributed significantly to the motivation and feasibility of this work.

---

**Supplemental Figure 1. ChIP-seq and ATAC-seq signal heatmaps.** (A) SWI/SNF occupancy sites from BIN67 cells (ChIP data re-analyzed from Pan *et al*. (14)) are annotated as proximal (overlapping TSS) or distal (>2kb) to TSS. Distal sites are separated by those overlapping SE sites (In SE), as determined by ROSE on H3K27ac ChIP data, and those non-overlapping those sites (Outside SE). Signal intensity over a 3kb region centered on the SWI/SNF peak center is shown with higher normalized read count in dark blue vs. white for absent read count. SMARCC1, DPF2, ARID2, and SMARCA4 ChIP-seq, and ChIP-seq Input enrichment are plotted on the same scale. Due to different read depth for H3K27ac ChIP-seq and ATAC-seq, these each had different scale maximums from the SWI/SNF subunits, but Control and SMARCA4 re-expression conditions are on the same relative scale for all samples. (B) ATAC-seq signal at SWI/SNF occupancy sites, categorized by change of H3K27ac at the same site with SMARCA4 re-expression. A cutoff of > |log_2_FoldChange in H3K27ac occupancy| was used to distinguish decreased/increased peaks from static peaks. Summary of average of peak intensity is shown to right of heatmaps, where dotted line represents SMARCA4 re-expression. (C) Total number of SWI/SNF occupancy peaks by location annotation (x-axis) and H3K27ac change after SMARCA4 re-expression (stacked bar chart). (D) Percent of SWI/SNF occupancy peaks by location annotation.

**Supplemental Figure 2. SCCOHT SE-associated oncogene expression compared to other ovarian cancers and normal samples.** In addition to Figure 3A, the remaining SE-associated oncogenes’ RNA expression levels. RNA expression from RNA-seq on SCCOHT tumors, age-matched TCGA high-grade serous ovarian carcinoma (HGSC) tumors, normal ovary tissue from ENCODE (ENCODE ovary), and ENCODE reference cell lines (ENCODE reference). FPKM + 0.01 is plotted on a logarithmic scale, with individual circles representing each sample and the box and whiskers plot summarizing the median value (heavy line), upper and lower 25% quartiles (box limits), and error bars showing highest and lowest values.

**Supplemental Figure 3. SALL4 expression in SCCOHT tumors.** SCCOHT TMAs stained for SALL4. Patient number is indicated to the left of the core images. Each tumor sample has at least one set of two representative cores from a given area, which are positioned next to each other in the row. Patients with more than one pair of cores are presented on separate rows. Staining score for each sample pair is indicated to the right.

**Supplemental Figure 4. Differential occupancy analysis within SCCOHT SEs.** Differential occupancy analysis was performed with csaw to identify peaks within BIN67 (A-B) and COV434 (C-D) SEs that change with triptolide treatment for 6 hours (A & C; N=4) and 16 hours (B & D, N=3). Cutoffs of | log_2_ fold change in peak signal | > 1.5 and false discovery rate of > 0.05 were used.

**Supplemental Figure 5. Triptolide and minnelide are well tolerated in mice bearing PDXs.** (A) Individual tumor weights of SCCOHT PDX-040 treated with vehicle (DMSO i.p.; N=10) or triptolide (0.6 mg/kg i.p., QD days 1-5, QOD days 10-60, N=4). (B) Individual tumor weights of SCCOHT PDX-465 treated with vehicle (PBS i.p., N=10) or minnelide (0.42 mg/kg i.p., QD, N=10). Individual mouse measurements shown in light lines. Group mean and standard error of the mean are shown in bold lines. (C) Individual body weights of mice bearing PDX-040 treated with triptolide. (D) Individual body weights of of mice bearing PDX-465 treated with minnelide. (E) SCCOHT PDX-040 and (F) PDX-465 tumor weights were equivalent between vehicle and treatment groups at study start.

