## Supplemental methods for "Landscape of super-enhancers in small cell carcinoma of the ovary, hypercalcemic type and efficacy of targeting with natural product triptolide"

*Genomic library preparations and sequencing*

CUT&RUN: DNA enriched in binding to K27-acetylated histone H3 (H3K27ac) was determined using CUT&RUN and sequencing. Cell lines were harvested at greater than 70% confluency with 0.05% trypsin. 10cm plates were seeded with 2-3x$10^6$ BIN67 or COV434 cells and allowed to adhere overnight. The next day, fresh media was added with 100nM triptolide with total DMSO concentration of less than 0.3%. After 6 or 16 hours of drug treatment, cells were harvested with 0.05% trypsin and total cell amount determined with cell count with dead cell exclusion with trypan blue staining using a Countess II. For tissues, we used 10-20mg frozen tissues and disassociated by pipetting in the first buffer in the protocol. EpiCypher CUTANA CUT&RUN protocol v1.5.2 was followed with 1x$10^6$ cells used for input of each cell line for both the H3K27ac and control (IgG) sample. H3K27ac antibody (Active Motif, cat#39133) was used at a 1:50 dilution (1ug/sample) in the antibody buffer and incubated overnight at 4°C. A normal rabbit IgG antibody (CST, cat#2729) was similarly used at a 1:50 dilution (1ug/sample). For targeted DNA cleavage, 2µL of EpiCypher CUTANA pAG-MNase (cat#15-1016) was added to each reaction. After completion of the CUT&RUN protocol, each sample was purified using the ChIP DNA Clean & Concentrator kit (ZYMO Research, cat#D5205). Samples were eluted in 30-50µL. Material was QC using an Agilent 4200 TapeStation with High Sensitivity D1000 ScreenTape (cat#5067-5584, reagents cat#5067-5585) and Qubit dsDNA HS Assay Kit (ThermoFisher, cat#Q32854). Typical yields were single digit nanograms with the main DNA fragment detected at ~150bp. KAPA HyperPrep Kit (Roche, cat#07962363001) was used to generate DNA libraries for sequencing. Protocol v6.17 was followed with following modifications: (a) 1ng of input DNA from the CUT&RUN experiments, (b) 0.5µL of 1.5µM KAPA UDI Adapter (Roche, cat#08861919702) adapter stock was used during adapter ligation step, (c) final KAPA bead purification used a first cut of 0.65x beads followed by

0.25x bead purification. Samples were QC as above using TapeStation and Qubit. Typical library yields were ~500ng. An equimolar pooling of libraries was performed for a final 1500pmols in 150µL with Tris pH8.5 buffer as final fill volume. Pooled libraries were loaded onto a NovaSeq6000 using a NovaSeq SP 100 cycle kit (cat#20027464) (51x9x9x51).

RNA-seq: Vehicle- or triptolide-treated SCCOHT cell lines were harvested in parallel to the CUT&RUN samples described above. RNA was extracted using Quick-DNA/RNA Miniprep Plus Kit (Zymo Research Cat#D7003), per manufacturer's instructions, with on-column DNase digestion. 500ng of total RNA was used to generate mRNA libraries using KAPA mRNA HyperPrep Kit library prep kit (Roche, cat#08098123702) per manufacturer's instructions. Final libraries were evaluated by TapeStation, quantitated using the Life Technologies/Invitrogen Qubit, and equimolarly pooled. Paired-end sequencing was performed on a NovaSeq6000 using a NovaSeq S4 300 cycle kit (cat#20028312) (151x11x11x151).

SCCOHT tumor RNA-seq was performed on a total of 10 tumors. Four tumors were previously described in Lang et al. (PMID: 29440177; dbGap accession phs001528.v1.p1, GEO accession GSE109919), which were sequenced to an average depth of 198.28 +/- 64.12 (standard deviation) million aligned reads. Six additional tumors were collected under IRB protocols: Western IRB protocol #1119451 at TGen (Samples TGEN-E through TGEN-I) and protocol #90-0573 at the University of North Carolina-Chapel Hill (Samples TGEN-J). This data is available at dbGaP (phs001528.v2.p1) and GEO (GSE216801). For the additional 6 tumors, RNA was harvested from formalin-fixed, paraffin-embedded (FFPE) SCCOHT tumors using the RNeasy FFPE Kit (Qiagen). Total RNA (200 ng) was heat fragmented on a GeneAmp PCR System 9700 (Applied Biosystems, Waltham, MA) to a target peak of 150 base pairs and libraries were generated using the SureSelect XT RNA (Agilent) capture following the manufacturer’s protocol. RNA libraries were quantified, and quality was evaluated using the Qubit DNA BR Reagent kit and Tapestation D1000 tapes. Libraries were pooled by equimolar ratios and sequenced on a HiSeq4000 (Illumina) using paired-end sequencing (77x9x77). Average sequencing depth was 77.05 +/- 26.44 (standard deviation) million aligned reads.

*Genomic analyses*

CUT&RUN read alignment and peak calling was performed using modified ChIP-seq workflows implemented in TGen's Phoenix pipeline (https://github.com/tgen/phoenix). First, BCL conversion was performed using Illumina's Bcl Converter tool and parsing by barcode into independent FASTQ files. Alignment to GRCh38 human reference genome was performed using bowtie2 (v2.4.2) (1) to generate BAM files (arguments: --local --very-sensitive-local --no-unal -- no-mixed --no-discordant --phred33 --minins 10 --maxins 700 -q), which were converted into CRAM format using samtools view -C. QC was assessed using FastQC (v0.11.8), samtools stats (v1.11) (2), and deepTools (v3.3.1) (3). Peaks were called using MACS2 callpeak (v2.2.6) (4) with the following arguments: -f BAMPE -B -q 0.01 --keep-dup all. After removal of peaks from the ENCODE Unified GRCh38 Exclusion LIst (blacklist; ENCFF356LFX), ROSE (5,6) and CREAM (7) were used to call SEs from the resulting narrowPeak files from MACS2. For ROSE, the ROSE_bedtoGFF.py was used to convert narrowPeak to GFF format, followed by ROSE_main.py (arguments: -g hg38 -s 12500 -5 2500). The R package CREAM tool was also used (arguments: MinLength = 1000, peakNumMin = 2). ROSE and CREAM SEs for each sample were concatenated together, and merged using bedtools. Genome regions were mapped to genes using ChIPpeakAnno (v3.24.2) and ROSE_geneMapper.py (ROSE v0.1). Oncogenes from the merged set of genes were identified from the COSMIC tier one cancer gene consensus list (v93, downloaded consensus table Jan 5, 2021). ChIPpeakAnno output was annotated with R package EnsDb.Hsapiens.v86 (v2.99.0) and ROSE_geneMapper.py output was annotated using the ROSE gene reference table (hg19_refseq.ucsc). Differential peak calling was performed using csaw (v1.30.1, windowCounts width=150, pe=”both”) with DESeq2 (v1.36.0), calling differential peaks within the SEs called from independent H3K27ac

CUT&RUN run on the same cell line. SE region intersections and upset plotting was performed with Intervene (v0.6.5,--bedtools-options f='.2').

Publicly available ChIP-seq data was also used to determine SWI/SNF binding at SEs and other genomic sites (GEO: GSE117735) (8). This analysis was performed from downloaded SRA files that were converted to FASTQs. Alignment and peak calling were performed as above with the following modifications/additions to MACS2 callpeak: -f BAM -q 0.001 --SPMR. SWI/SNF peaks were defined by using bedtools intersect to find overlap of SMARCC1 with DPF2 or ARID2 for BAF and PBAF peaks, respectively, which was performed separately on BIN67 control and BIN67 SMARCA4 WT re-expression. All BAF and PBAF sites in control and SMARCA4 re-expression conditions were merged for the full set of SWI/SNF binding sites. IP enrichment at each of these sites was calculated using bedtools coverage with the resulting BED file of SWI/SNF binding sites as -a, and SMARCC1, H3K27ac, and Input in control and SMARCA4 WT re-expression BAM files as -b. The read counts were then normalized as follows: (total IP signal in region per million mapped reads + pseudocount of 0.1)/(total input signal in region per million mapped reads + pseudocount of 0.1). Annotation of SWI/SNF binding sites was performed using ChIPPeakAnno (v3.22.4) (9,10) with built-in hg38 transcription start sites (data("TSS.human.GRCh38") and featureType = "TSS"). Proximal sites were defined as those that overlapped with TSSs, and Distal sites were those >2000kb from nearest TSS. Distal sites were further assessed for falling within or outside SEs called from ROSE from the same dataset (merged from H3K27ac-derived SEs from control and SMARCA4 re-expression). Heatmaps were generated using deepTools with normalization with the RPKM method.

RNA-seq data was analyzed using RNA workflows implemented in TGen's Phoenix pipeline (https://github.com/tgen/phoenix). Briefly, alignment of FASTQ files against the GRCh38 human reference genome was performed using using STAR (v2.7.7a) (11), and gene expression estimates were performed using Salmon (v1.4.0) (12). Gene-level quantification in

Transcripts Per Million kilobases (TPM) were used in subsequent steps. Where separate batches were analyzed together (as in cell line and tumor RNA-seq heatmaps), batch correction was performed by first performing variance-stabilizing transformation, then using removeBatchEffect in the limma R package (v3.44.3) (13). Differential expression analysis was performed using DESeq2 (v1.30.0) (14), using cutoffs for $log_2$FoldChange of +/- 1.5 and adjusted p-value < 0.05 to determine significance. Heatmaps were generated using pheatmap in R using 'complete' clustering method and Euclidean distances. For input to gene ontology analysis, SE-associated genes in the top 2 high expressing clusters for gene expression (Fig. 2C, purple box; Supplemental Table 1) were input into PANTHER v16.0 (http://geneontology.org/), and terms with FDR ≤ 0.05 were visualized. For comparison of SCCOHT data to TCGA and ENCODE normal ovary and reference cell lines, FPKM expression values were used.
