## Supplemental Figures for "Landscape of super-enhancers in small cell carcinoma of the ovary, hypercalcemic type and efficacy of targeting with natural product triptolide"

### Supplemental Figure 1.

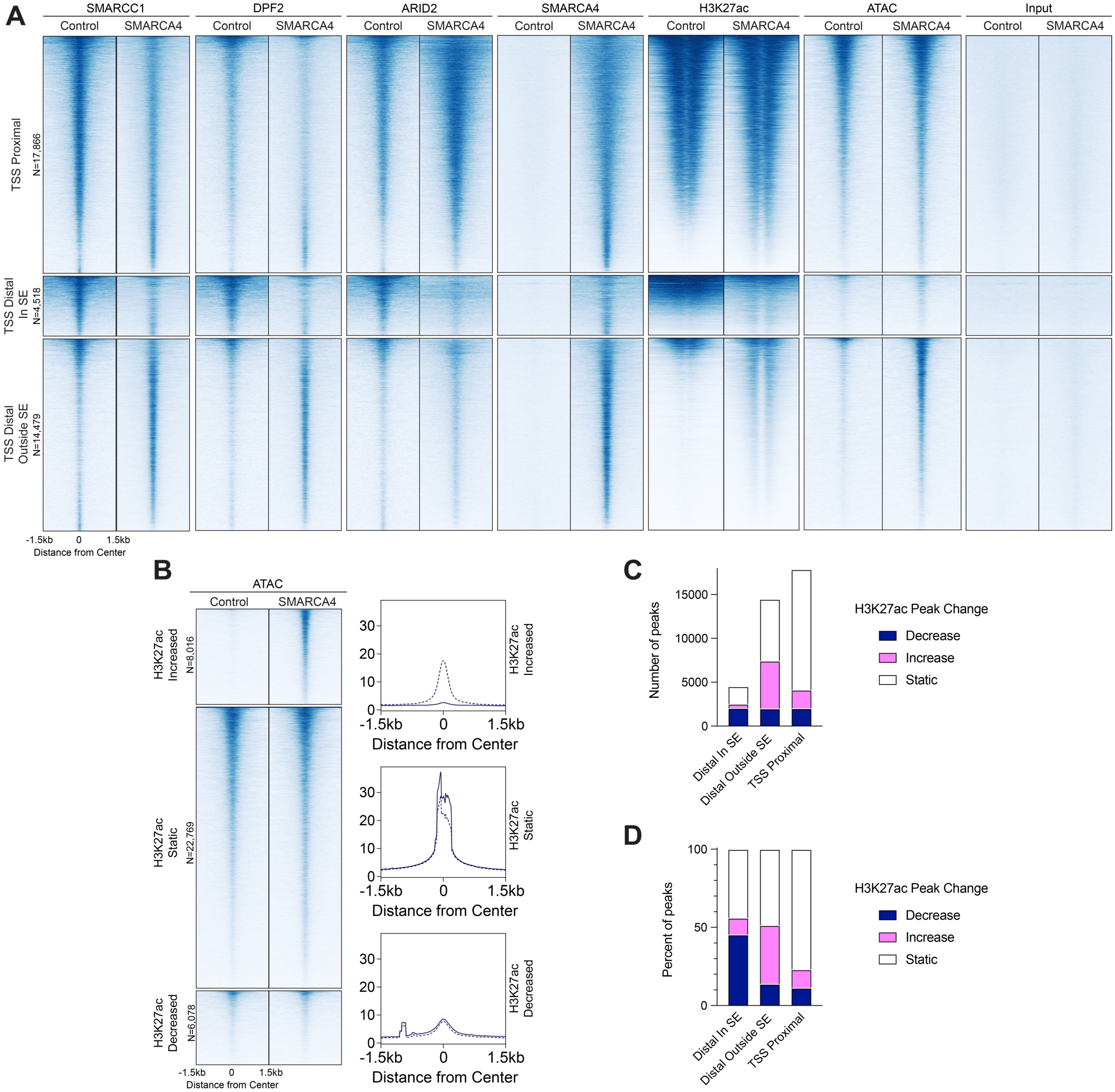

### Supplemental Figure 2.

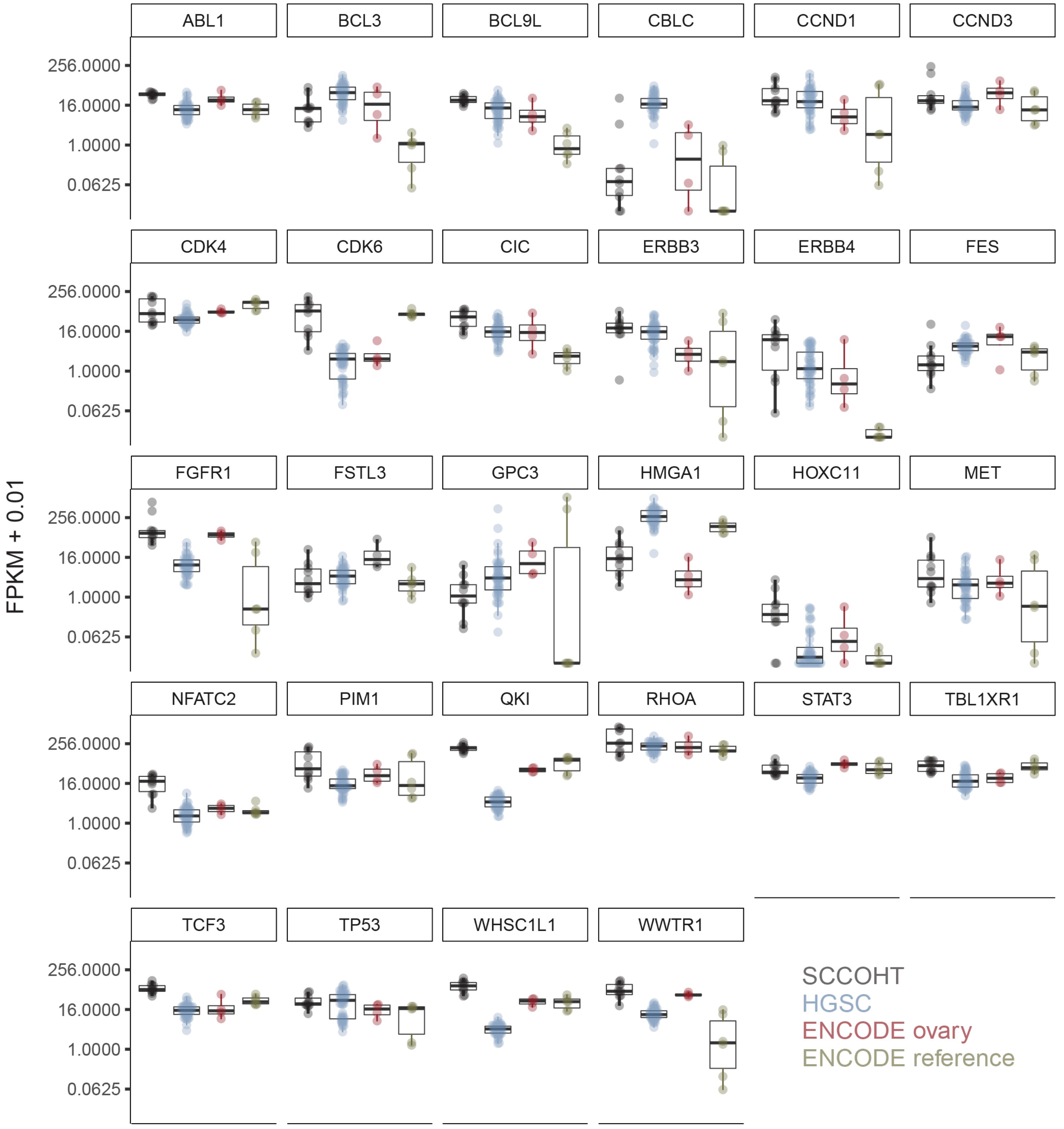

### Supplemental Figure 3.

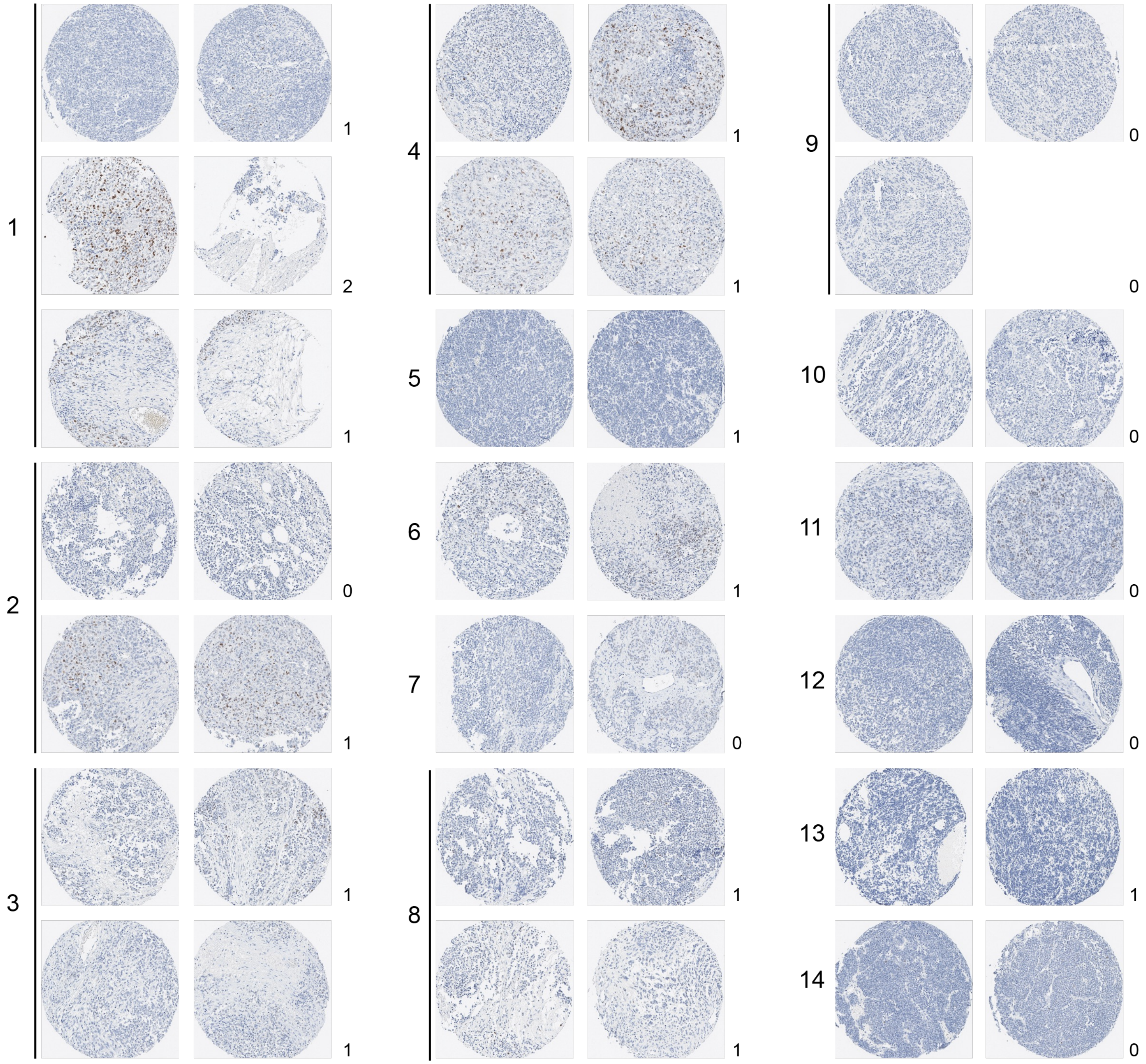

### Supplemental Figure 4.

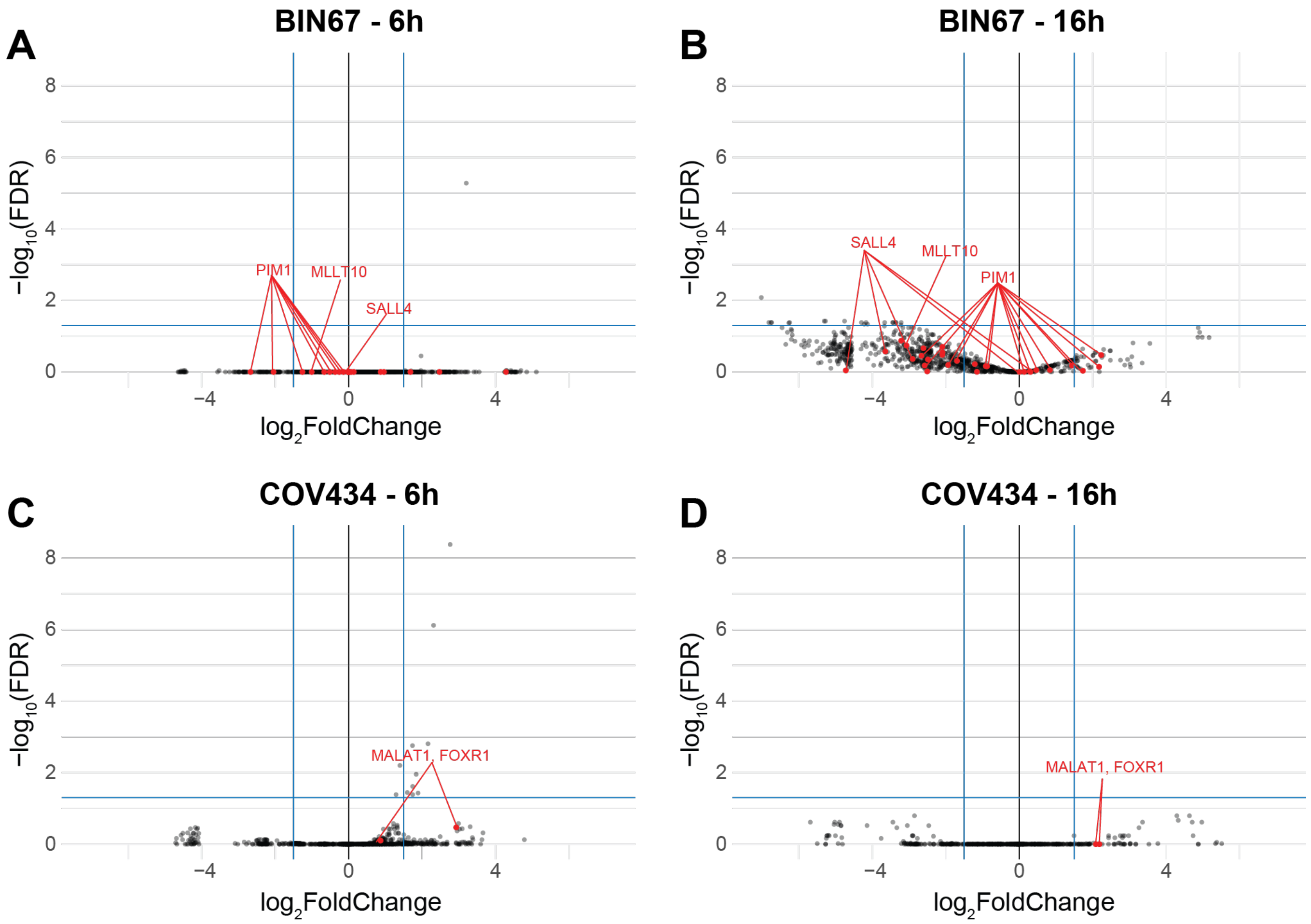

### Supplemental Figure 5.

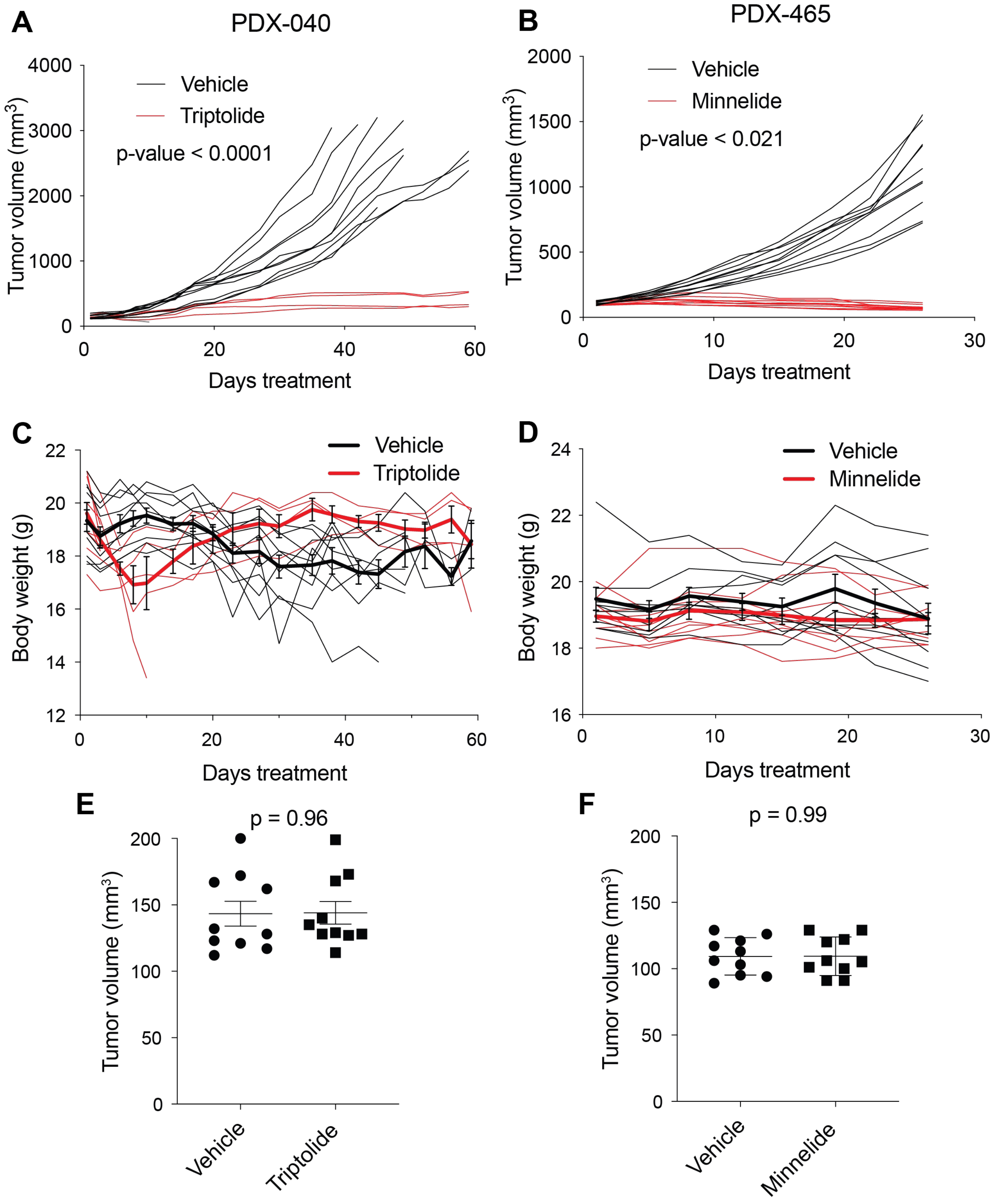
